# TiDeTree: A Bayesian phylogenetic framework to estimate single-cell trees and population dynamic parameters from genetic lineage tracing data

**DOI:** 10.1101/2022.02.14.480422

**Authors:** Sophie Seidel, Tanja Stadler

## Abstract

The development of a multicellular organism is governed by an elaborate balance between cell division, death, and differentiation. These core developmental processes can be quantified from single-cell phylogenies. Here we present TiDeTree, a Bayesian phylogenetic framework for inference of time-scaled single-cell phylogenies and population dynamic parameters such as cell division, death, and differentiation rates from genetic lineage tracing data. We show that the performance of TiDeTree can be improved by incorporating multiple sources of additional independent information into the inference. Finally, we apply TiDeTree to a lineage tracing dataset to estimate time-scaled phylogenies, cell division, and apoptosis rates. We envision TiDeTree to find wide application in single-cell lineage tracing data analysis which will improve our understanding of cellular processes during development. The source code of TiDeTree is publicly available at https://github.com/seidels/tidetree.

## Introduction

Understanding the principles of development is a major goal for developmental, regenerative and cancer biology. Cell phylogenies contain rich information on cellular events during development; they depict the parental relationships between cells, map the origin of cell types and contain a signal for key developmental parameters, such as the cell division, death and differentiation rates [27]. Recent developments in genetic lineage tracing provide the data to reconstruct such cell phylogenies and use them to quantify developmental processes.

Several genetic lineage tracing systems, or recorders, have been developed [22, 24, 1, 26, 18, 15], all relying on an enzyme, such as CRISPR-Cas9, to edit, or scar, genomic target regions that are passed on to successive generations. Hence, they provide a record of the ancestral relationships between cells and can be used to reconstruct a cell phylogeny.

Several computational methods have been developed to reconstruct cell phylognies from lineage tracing data. Initially, methods based on maximum parsimony [22] were used and custom algorithms for cell phylogenies were developed [26]. These methods aim to reconstruct a tree that minimises the number of edit acquisition events. However, the assumption that edit acquisition is rare is violated by recurrent editing outcomes of the CRISPR-Cas9 enzyme [21]. To reduce biased inference, frequently occurring scars had to be excluded during data pre-processing, resulting in data loss [26].

More recently, methods were developed that can incorporate *a-priori* information on the frequency of editing outcomes [16, 14, 13], which can help to reduce biased inference due to homoplasy. While some approaches focus on improving scalability to reconstruct trees with millions of cells [16, 19], others focus on detailed modelling of the editing process to enable more accurate inference [13]. A key example of the latter is the maximum likelihood framework GAPML [13], which models the editing process of a GESTALT recorder [22]. Additionally, a molecular clock assumption is employed allowing to order cell division events (branching events in the phylogeny) relative to each other.

Nevertheless, until now no framework exists that can infer time-scaled trees, that is phylogenies where branch lengths are scaled in absolute duration and a time is associated with each cell division. A time-scaled tree provides a temporal scaffold along which one can subsequently integrate other multi-modal single-cell measurements. Additionally, all existing methods only provide a single best estimate of the tree linking cells together and ignore the phylogenetic uncertainty in this estimate.

To overcome these limitations, we developed a time-dependent editing model and show how to calculate its likelihood. We implemented this model in TiDe-Tree (Time-scaled Developmental Trees), a package within the BEAST2 [6] platform. We show how to use it for Bayesian phylogenetic inference of time-scaled cell lineage trees, which enables the inference of phylodynamic parameters alongside phylogenies and further provides a natural framework for quantifying uncertainty and incorporating prior information. We show how integrating commonly available additional information can result in more accurate and precise estimates. Finally, we apply TiDeTree to lineage tracing data [8] and estimate time-scaled trees and phylodynamic parameters. Compared to other methods [14] TiDeTree’s ability to recover the correct tree topology is always among the top three, while also being the only method to estimate cell division times and to quantify uncertainty.

## Materials and Methods

### Phylogenetic model for lineage tracing

Here, we introduce a time-dependent editing model. Based on this model, we derive the likelihood function to perform phylogenetic inference from lineage tracing data.

#### A general lineage recorder

We consider the following setup of a lineage recorder: A precursor cell contains *m* genomic regions, henceforth called *targets*, that are targeted by an editing enzyme. Different targets can be distinguished by target-specific *barcodes*. We refer to the combined region of a target and its barcode as an *integration*.

Given such a setup, the experiment starts with a precursor cell where all targets are unedited (depicted as state 0 in Fig. 1). During a time period from *t*_1_ to *t*_2_, the *scarring window*, any target can be scarred, i.e., transition from the unedited state (0) to one of *S* scarred states (e.g., state 2 in Fig. 1). Experimentally, this is implemented either via injection of the editing enzyme into the precursor cell (*t*_1_ = 0) or by inducing the enzyme’s expression later during development (*t*_1_ *>* 0). Usually, *t*_2_ is determined by independent experiments that identify at what time point the fraction of unedited integrations stops decreasing [1, 26]. Throughout the experiment a target can be silenced at any time (state 1 in Fig. 1).

**Figure 1:**
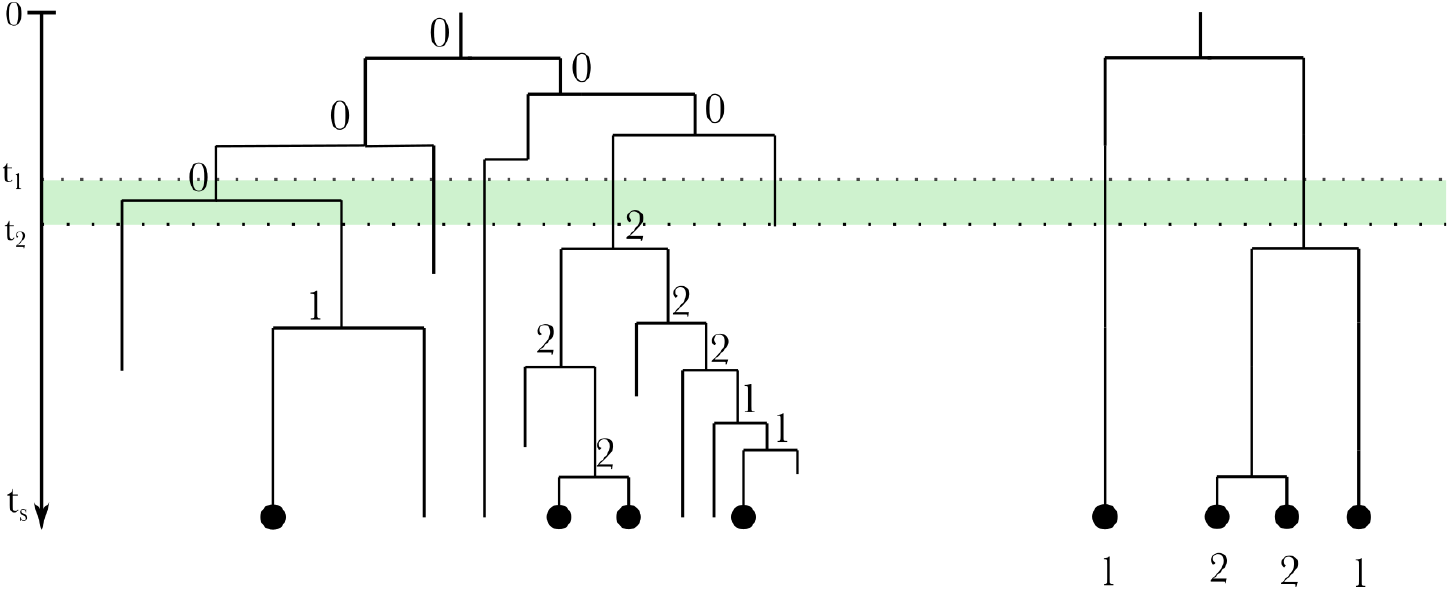
Example cell phylogeny for initial cell with one integration. Numbers denote the target state at each node: 0 corresponds to an unedited, 1 to a silenced and 2 to a scarred target. Targets can be silenced at any time. Scarring can only occur during the scarring window (green shading). Internal nodes represent cell divisions and branch lengths the time between them. Left: Complete developmental phylogeny with all cells and their target state at internal nodes. Right: Reconstructed phylogeny of only the cells sampled at time *t*_*s*_.

At the end of the experiment, at time *t*_*s*_, a subset of cells is selected for sequencing to determine the states of their integrations. Either DNA or RNA sequencing may be used, as long as the readout allows to assign integrations to a single cell. Both sequencing technologies might fail to report the presence of an integration due to dropouts during the sequencing process. Additionally, RNA sequencing cannot record an integration if the genomic region containing the integration was silenced during development.

#### Substitution model

We now formulate a model for the lineage recorder introduced in the last section. We model barcode evolution as a continuous time Markov chain with state space Ω = {unedited, silenced, scarred}, initial state *X*_0_ = {unedited} and a timedependent (piecewise-constant) transition rate matrix:

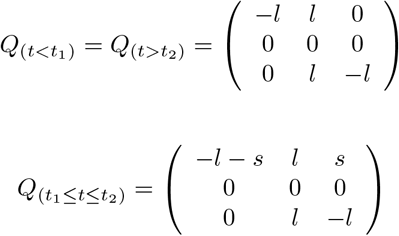

Silencing of a barcode’s genomic region occurs at constant rate *l* throughout the experimental period, while scarring only occurs during the scarring window, *t*_1_ ≤ *t* ≤ *t*_2_, at constant rate *s*.

Note that scarring and silencing cannot be reversed. Further, while both unedited and scarred barcodes may be silenced at any time, once silenced a barcode cannot be scarred, and once scarred cannot be scarred again.

In the following we extend the simple substitution model above to *S* scarred states. Additionally, we introduce the global editing rate *r* at which site *S* accumulates a scar per unit of time (*r* corresponds to the molecular clock rate in traditional phylogenetics). We arbitrarily set *s*_*S*_ = 1 and generally assume that site *i* accumulates scars at rate *r* × *s*_*i*_. Thus one can interpret the scarringrates *s*_*i*_ as the rates at which particular scarred states *i* arise relative to state *S*.

In our substitution model, this leads to increasing the the state space of the Markov chain to Ω = {unedited, silenced, scar_1_, scar_2_, …, scar_*S*_}:

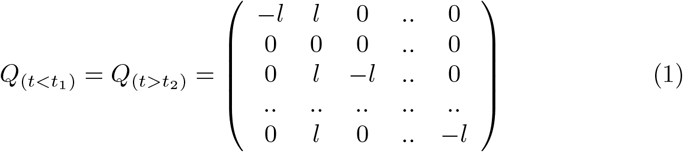

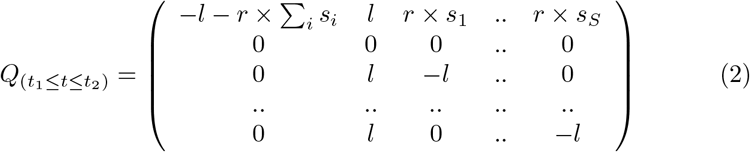

Note that this substitution model is not time reversible. Generally, time reversibility is a desirable property, because it ensures that the transition rate matrix can be diagonalised, which simplifies computation. However, the transition matrices (Eq. 1, 2) are diagonalisable and the transition probability matrix, *P*_*t*_, can be determined analytically (see Appendix). As a result, we can compute *P*_*t*_ in *O*(*k*), when otherwise matrix diagonalisation would require *O*(*k*^3^), where *k* = *S* + 2 is the dimension of the rate matrix.

In practice some scarred states may occur more often than others. Especially frequently occurring scars can bias phylogenetic inference (due to homoplasy). By allowing for scar state-specific scarring rates, *s*_*i*_, we avoid this bias. Essentially, this allows us to weigh the information content a scarred state provides since, for example, a scarred state with a high scarring rate is likely to arise several times on the tree, while a scarred state with low scarring rate will likely arise only once.

#### Phylogenetic likelihood

To calculate the likelihood of the model parameters given the data (target states of the sampled cells) and the model, we introduce the following notation. Let *T* − be a tree with *n* tips representing the reconstructed phylogeny of the sampled cells. This tree has *n* tip nodes of degree 1, *n* − 1 internal nodes of degree 3, including the root node, the most recent common ancestor of all samples. Additionally, we have an origin node of degree 1 ancestral to the root node which specifies the start of the experiment (Fig. 1 right). By convention, we number internal nodes from (*n* + 1) to (2*n*) from the tips toward the origin. We further subdivide all branches at time points *t*_1_ and *t*_2_ and label these additional nodes of degree 2 with 2*n* + 1, 2*n* + 2, … 2*n* + *d* + 1. Further, let *τ*_*i*_ be the length of the branch that connects node *i* to its parent, *π*_*i*_. ∈

Let *θ* summarise the parameters of the transition rate matrix, i.e., the scarring rates *s*_1_, …, *s*_*S*_, clock rate *r*, and silencing rate *l*. Note that each branch is associated with one transition rate matrix (matrix in Eq. 1 or 2), i.e. the process does not change along a branch. We use vector *b*_*i*_ to refer to the state of all integrations in node *i* and specify with *b*_*i,j*_ the state of the *j*^th^ integration for *j* 1, …, *m*. For the tips, these *b*_1_, …, *b*_*n*_ are known. Then, we can calculate the likelihood of the tree and parameters *θ*, given the target states of all *m* integrations at the sampled cells (*b*_1_, …, *b*_*n*_) by summing over all internal node states:

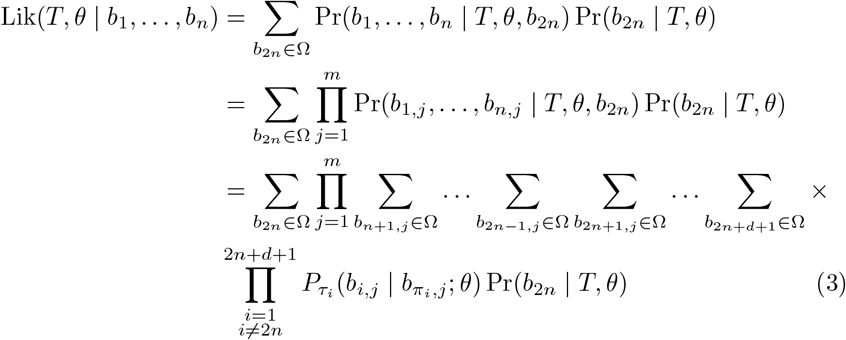

where 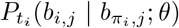 represent the transition probabilities from parent node *π*_*i*_ to its child node *i* along branch length *t*_*i*_. They are derived from the transition probability matrices (Appendix, Eq. 9 and Eq. 10).

All ancestral targets, i.e. targets at the origin, are unedited (by the experimental design). Thus, the probability of the origin state being unedited (0) is

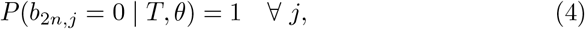

We implemented the substitution model and the likelihood calculation employing the pruning algorithm [12] within the package Time-scaled Developmental Trees (TiDeTree) to be used within the widely used BEAST2 [6] platform, thus enabling phylogenetic and phylodynamic inference under the model described above.

### *In-silico* validation of the implementation

We validate TiDeTree on synthetic data using well-calibrated simulations [9]. We simulate a tree under a birth-death-sampling process [28], mimicking the growth process of tissue from a single cell. After 32 time units, we use a sampling probability *ρ* to draw sampled lineages uniformly at random from the surviving lineages, conditioning on sampling at least one lineage. This step models the random sequencing of a subset of cells from the entire tissue and results in the sampled tree.

To obtain an alignment, we simulate the evolution of 10 targets from the origin (the start of the process at *t* = 0) along the sampled tree to each of the contemporaneous tips under the editing model summarized in Eq. 1 and 2. The result is an alignment, where we know the target state of each cell. From this alignment we seek to infer the phylogeny, together with the parameters that were used in the simulation.

We apply the above simulation procedure 1000 times. For every simulation, we draw all parameters but the death rate from their respective prior distributions (see Tab. 1). The death rate is fixed to 0.4 for two reasons. Firstly, it helps to get a distribution of simulated trees that is neither too large, nor too small. Secondly, when using the birth-death model for inference, one of its parameters has to be fixed for identifiability reasons [28]. We further fix one of the scarring rates, namely we set *s*_50_ = 1. This allows us to interpret all other scarring rates relative to *s*_50_. We discard trees with more than 700 sampled lineages which constitute 14% of the data such that the final dataset consists of 860 alignments.

**Table 1:** Prior distributions for *in silico* validation.

| Parameter | Prior | Percentiles: [5<sup>th</sup>, 95<sup>th</sup>] |
| --- | --- | --- |
| scarring rates | Exponential(2) | [0.03, 1.5] |
| clock rate | LogNormal(-7, 0.5) | $[4 \times 10^{-4}, 2 \times 10^{-3}]$ |
| birth rate | LogNormal(-0.6, 0.1) | [0.47, 0.65] |
| sampling proportion | Beta(4, 8) | [0.14, 0.56] |

For inference, we apply TiDeTree to each input alignment and specify the same prior distributions as before (Tab. 1). We again keep the death rate and scarring rate, *s*_50_, fixed at their respective true values. Further, we keep the origin constant at 32, which incorporates our knowledge of the duration of the experiment. Additionally, we fix the duration of the scarring process to the truth (i.e., 0 to 16 time units). This information is readily available in existing studies. We let the analysis run for 10^8^ steps or until the effective sample size (ESS) is above 200 and discard 10% of the analysis to account for burn-in.

We compute the coverage of all parameters as the fraction of times the true parameter was contained in the 95% highest posterior density (HPD) interval. Further, we report the Pearson correlation coefficient between the estimated posterior medians and true values.

### Assessing accuracy and precision of parameter inference based on our simulations

Next, we explore how accurate and precise the parameters of our model can be inferred from simulated data. For that we make different assumptions for the prior distributions and available data for the inference.

As before, we simulate 1000 trees and alignments, except that we do not draw the parameters from a prior distribution but use a fixed value for each parameter (Tab. 2 and Supplementary Tab. S2).

**Table 2:**
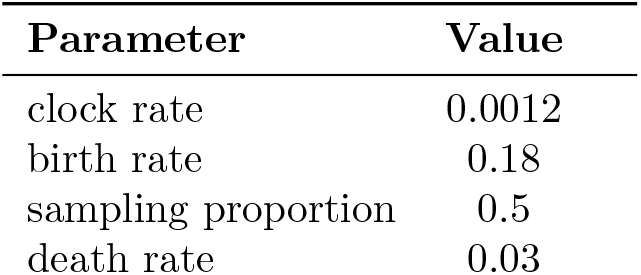
Parameters for simulation study.

| Parameter | Value |
| --- | --- |
| clock rate | 0.0012 |
| birth rate | 0.18 |
| sampling proportion | 0.5 |
| death rate | 0.03 |

#### Weakly informative prior distributions

First, we assume that we have limited information on all parameters that we use as priors in our analysis (Tab. 3). We fix the sampling proportion and the global editing rate to their true values. Then we apply TideTree to the first 100 alignments to infer the phylogeny, and scarring, birth and death rates. We let the analysis run for 24 h or until the effective sample size is above 200 and discard 10% of the analysis to account for burn-in.

**Table 3:**
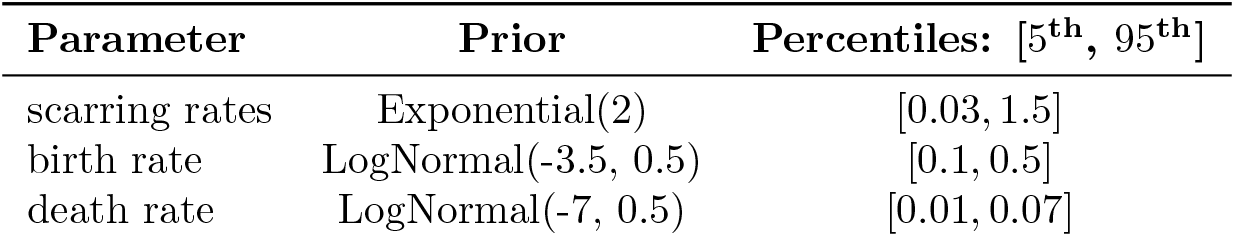
Prior distributions for simulation study.

#### Relative scarring rate information known

In experimental studies, information on the relative occurrence of the scarring outcomes can easily be obtained. For example, scarring can be induced in a population of cells and the relative frequency of the resulting scars can be determined by bulk sequencing. This information can be inserted into the inference by setting the relative scarring rates to the relative scarring frequencies.

To showcase parameter inference given this type of information, we mimic such a scenario in a simulation study. We reuse the first 100 alignments from the previous scenario. Then, we perform inference as before, but fix all relative scarring rates *s*_*i*_ to the truth during the inference.

#### Relative scarring rates known and experimental replicates

Another source of information comes from experimental replicates. The same experiment is often performed several times and hence yields to multiple draws from the same stochastic developmental process. In such a scenario, the relative scarring rates, the birth rate and death rate are identical across replicates.

To showcase parameter inference for multiple replicates, in addition to known scarring rates, we use all 1000 alignments from the simulated data. We distribute the alignments in such a way, that there are 10 input alignments for each inference run (mimicking 10 experimental replicates). In particular, we enrich each of the 100 alignments above by another 9 alignments from the set of 1000 alignments. Additionally, we infer a tree for each alignment and share the birth rate and death rate across trees.

#### Metrics to assess accuracy and precision of inferred parameters

To evaluate the performance of TiDeTree with respect to the correct tree topology, we do not use popular metrics such as the Robinson-Foulds distance, because we can only evaluate the correct topology for internal nodes of height larger than 16. As editing stops after 16 time units, there is no signal to differentiate between tree topologies for nodes below that height. Hence, we compute the mean and minimum posterior probability of all true internal nodes with height larger than 16 for each tree (that is, all nodes that fall within the scarring interval). For each scenario, we compute the average mean and minimum posterior probability by averaging across 100 trees. For the third scenario, i.e. where scarring rates and 10 experimental replicates are provided, we compute this metric using only the first tree that is identical to those provided in the other scenarios.

The remaining phylogenetic parameters, tree height and tree length, as well as the birth rate and the sampling proportion are evaluated based on the average bias, root mean squared error (RMSE), HPD width and coverage. The HPD width is the difference between the upper and the lower bound of the 95% posterior interval normalised by the true value of the parameter. The average bias and RMSE are also normalised by their respective true parameters.

### Application to experimental lineage tracing data

We apply TiDeTree to data used in the DREAM challenge [14], generated by the intMEMOIR system [8], where the ground truth cell phylogenies are known. In the analysis we quantify the ability of TiDeTree to recover the true tree topology, tree height, tree length and phylodynamic parameters.

#### Experimental setup

The experiment consists of mouse embryonic stem cells replicating for 54 hours. Each starting stem cell carries an intMEMOIR recording array with 10 unedited targets. During the first 36 hours, the expression of an integrase is induced, that can either invert or delete a target. Once the target is edited, it cannot be edited again. During the whole experimental period, cells are tracked using time-lapse microscopy. Therefore, the true topology of the cell phylogeny is known. As the movie frames are taken approximately every 15 minutes, we can only pinpoint the time of a cell division event within a 15 minute interval around the truth.

#### Testing for temporal signal in the data

In phylogenetics the concept of a molecular clock encapsulates the assumption that the expected number of edits accumulating over time is constant. Under this assumption the tree’s branch lengths in number of edits can be converted to calendar time.

We checked whether the molecular clock assumption holds for the intMEMOIR dataset. We used the true trees and the target states at their tips and calculated the expected number of edits at each internal node. Basically, we used a parsimony argument to compute each state of the parent *c*_parent_ given its children:

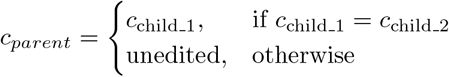

This gives us a lower bound on the number of edits that occurred on each branch and an expected number of edits until every internal node in the true tree.

#### Phylodynamic inference

##### Unsupervised inference

When performing lineage tracing using a lineage recorder, the true trees are not known. Thus there is no ground-truth information which might be used as a training set. Thus, to evaluate the performance of our method, we perform phylogenetic inference without using the information from the ground truth trees.

We assume that the editing process is governed by the product of two rates. Firstly, the “clock rate” *r* captures the rate that a deletion occurs per unit of time. Then, we introduce the inversion rates relative to *r*. We assume that inversions occur at rate 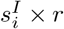. As no silencing occurs, we set the silencing rate to 0. Hence, we have the following site dependent transition matrix that derives from Eq. 2:

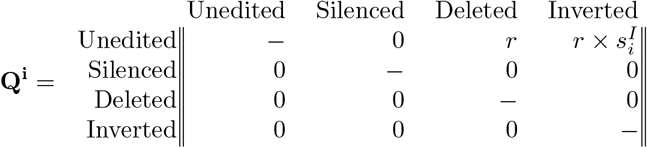

We incorporate our *a priori* knowledge on the editing process in priors on the editing parameters (see Tab. 4). These encapsulate our *a priori* belief that between 1 and 10 edits should occur on a recording array over the time course of 54 h. We place a weakly informative prior centered on 1 on the relative inversion rate, stating that we expect deletions and inversions to appear with similar frequency.

**Table 4:** Prior distributions for intMEMOIR analysis.

| Parameter | Prior | Percentiles: [5<sup>th</sup>, 95<sup>th</sup>] |
| --- | --- | --- |
| clock rate $r$ | LogN(-5, 1) | [0.001, 0.03] |
| inversion rate $s_I^i$ | LogN(0, 1) | [0.2, 5.2] |
| growth rate | Uniform(0, 0.1) | [0.005, 0.095] |
| turnover | Uniform(0, 1) | [0.05, 0.95] |
| scarring start | Exp(2) | [0.026, 1.5] |

We again use a birth-death-sampling model as the phylodynamic model to infer cell division and death rates. Given the small size of the cell population (4 − 40 cells), we expect the population growth process to be highly stochastic. The birth-death-sampling model can account for these stochastic fluctuations. We specify a prior on the growth rate (birth rate −death rate) stating that we expect less than 220 cells at the end of the experiment (after 54 h). Additionally, we place a prior on the cell turnover (death rate */* birth rate), stating that we expect the birth rate to be larger than the death rate (Tab. 4). We fix the sampling proportion at present to the ratio of cells in the alignment over the total number of cells in the colony (e.g. from [8], Supplementary Table 2.)

We run 5 independent MCMC chains for 24 h, resulting in ∼9×10^7^ iterations per chain and assess their convergence using Tracer [25].

##### Using independent prior information

In [8], the authors published data on the editing outcomes of a large number of barcodes after 36 h. Using this data, we can extract information on the relative occurrence of inversion over deletion and specify an informative prior. We use bootstrapping to approximate the distribution of inversions over deletions for each site and use this as prior information in the inference. Specifically, for each site, we bootstrap the ratio of inversions over deletions using 10^4^ bootstrap replicates. From the bootstrap samples, we compute the median, 5^th^ and 95^th^ percentiles and then specify a log-normal prior that is similar in the percentiles. For the exact specification see Tab. 5.

**Table 5:** Ratio of inversion over deletion per site assumed in the inference. The estimated percentiles are obtained using bootstrapping of the data in [8]. Lognormal prior parameters *µ* and *σ* are obtained based on the estimated percentiles.

| Site | Percentiles: [5<sup>th</sup>, 50<sup>th</sup>, 95<sup>th</sup>] | $\mu$ | $\sigma$ |
| --- | --- | --- | --- |
| 1 | [0.53, 0.59, 0.65] | -0.55 | 0.1 |
| 2 | [1.07, 1.35, 1.7] | 0.3 | 0.2 |
| 3 | [1.56, 1.79, 2.05] | 0.6 | 0.1 |
| 4 | [0.7, 0.76, 0.84] | -0.3 | 0.1 |
| 5 | [1.09, 1.23, 1.38] | 0.2 | 0.1 |
| 6 | [15.5, 21.57, 32.6] | 3.1 | 0.3 |
| 7 | [0.76, 0.89, 1.04] | -0.1 | 0.11 |
| 8 | [0.9, 1, 1.1] | 0 | 0.1 |
| 9 | [0.94, 1.04, 1.14] | 0 | 0.1 |
| 10 | [6.56, 8.75, 12.17] | 2.2 | 0.2 |

As explained above, editing is induced from the start of the experiment. However, editing may actually only start after a delay, as suggested based on exploratory analysis of the intMEMOIR dataset (see Results). We therefore treat the editing interval starting time as a parameter that we sample during the inference. We place an exponential prior with mean 2 on it, to express our expectation that scarring should start shortly after the onset of the experiment.

#### Comparison of results using TiDeTree to results from the DREAM challenge

For each alignment, we infer a tree posterior distribution using TiDeTree. For every posterior, we generate the maximum clade credibility (MCC) tree using TreeAnnotator [6] with default settings. Then, we compute the normalised triplet and Robinson-Foulds distance between each MCC tree and its corresponding true tree using TreeCmp [4] as done in the DREAM challenge and average over all datasets. We use the benchmarked set of competitor methods from the DREAM challenge and compare our unsupervised inference against them.

## Results

We developed a time-dependent editing model and derived its likelihood function, that enables phylogenetic inference from genetic lineage tracing data in a Bayesian Markov Chain Monte Carlo (MCMC) framework. We implemented these calculations as a BEAST2 [6] package called Time-scaled Developmental Trees (TiDeTree), enabling the inference of phylodynamic parameters alongside the phylogenetic trees. We validate TiDeTree using well-calibrated simulations and show different scenarios for integrating additional information. Finally, we apply it to a publically available lineage tracing dataset, which contains data from mouse embryonic stem cells as well as the ground truth trees. We estimate the time-scaled developmental phylogenies, cell division and death rates.

### *In-silico* validation of the implementation

We validate TiDeTree using well-calibrated simulations [9]. For the inferred parameters in the simulation study, we compute the coverage, i.e. the number of times the true parameter was contained within the 95% posterior interval, for each parameter. The coverages converge to 95% for all inferred parameters (Supp Tab. S1) indicating correctness of the implementation.

### Assessing accuracy and precision of parameter inference based on simulated data

To assess the TiDeTree’s capacity to correctly infer parameters from the data, we contrast the true values against the estimated medians in Fig. 2. TiDeTree infers the tree height (time between the most recent common ancestor of all cells and the time of sampling) well for all heights above 16 (green shaded area in Fig 2A). For heights below 16, the posterior HPD intervals are wider (mean relative HPD width of 8.7 compared with 0.4) and the posterior medians do not correlate as well with the true values (R=0.71, CI=[0.61, 0.79] compared with R=0.81, CI=[0.78, 0.83]). This is to be expected given our simulations, as no new edits are recorded after 16 time units and thus there is no signal to inform the tree height. The tree length (sum of all branch lengths) and the birth rate are recovered and correlated (with R=0.987, CI=[0.985, 0.989] and R=0.69, CI=[0.66, 0.73], respectively). There is little signal to inform the sampling proportion as posterior HPD intervals are broad and posterior medians are only weakly correlated to the truth (R=0.43, CI=[0.37, 0.48]).

**Figure 2:**
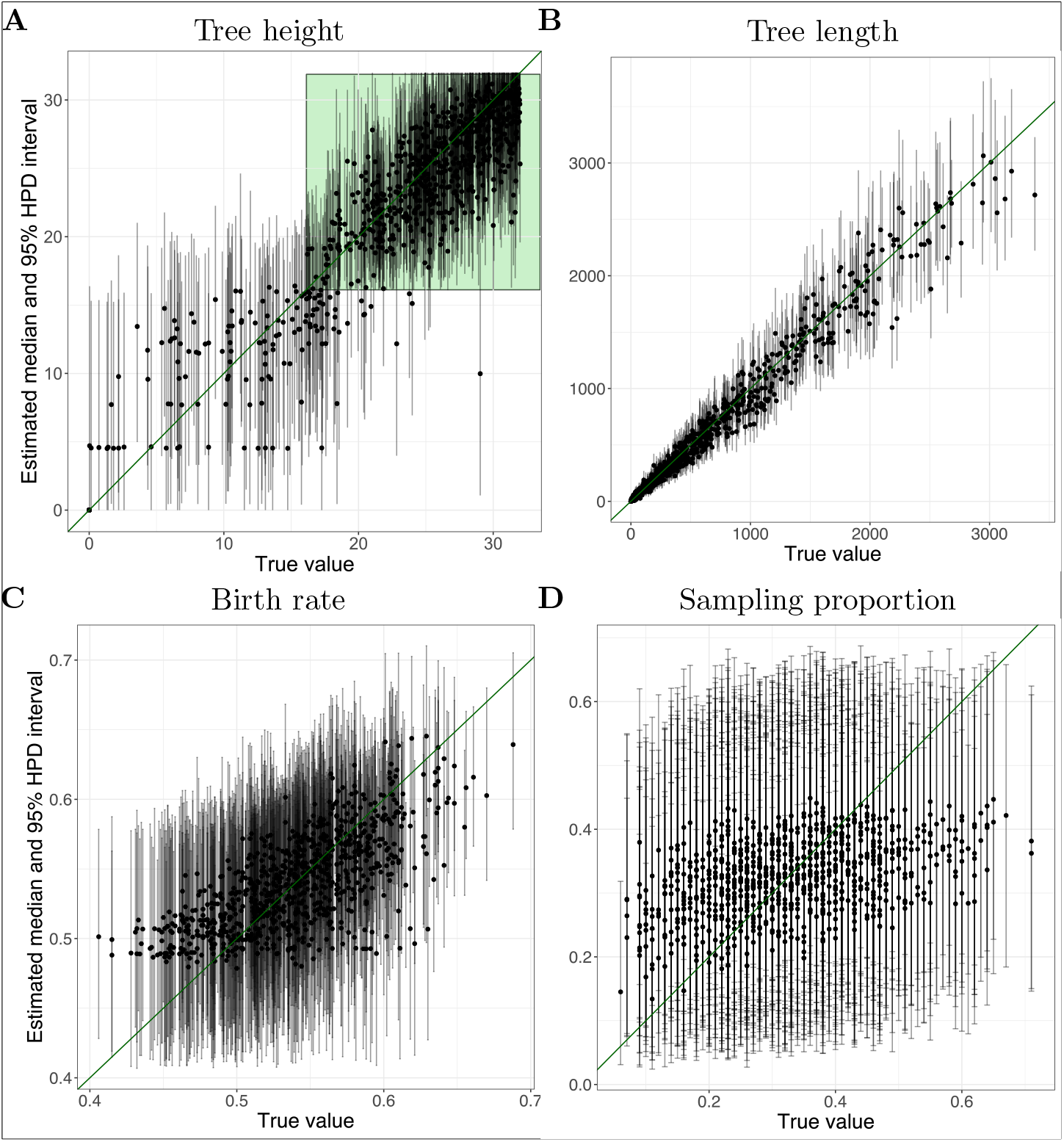
Parameter estimates based on simulated data. The graphs show the median estimates (black dots) and 95% HPD intervals (grey lines) on the y-axis and the true values on the x-axis based on 860 simulations. Four different parameters are shown, the (A) tree height, (B) tree length, (C) birth rate and (D) sampling proportion. The diagonal green line indicates the performance of a perfect estimator. The green shaded area in panel A illustrates the time period in which editing takes place.

### Assessing accuracy and precision of parameter inference when integrating additional information

In a second simulation study, we simulate 1000 alignments. With these, we show how commonly available independent information can improve the inference. As our baseline, we apply TiDeTree to infer the tree, scarring rates, birth and death rate from the first 100 alignments, using one alignment per inference and weakly informative priors as input.

The relative frequencies of different scars are often known from an independent experiment, e.g., by letting a population of cells be edited for a short time period and then performing bulk sequencing. If not, several studies [2, 7, 23] have probed the sequence dependence of the scarring outcomes of CRISPR-Cas9 and allow to estimate relative frequencies. Such information can be used to fix the relative scarring rates, or to set prior distributions on them. To mimic this scenario, we use the same first 100 alignments and perform inference as in the baseline, but also fix the scarring rates to their true values (Fig. 3A, scenario B).

**Figure 3:**
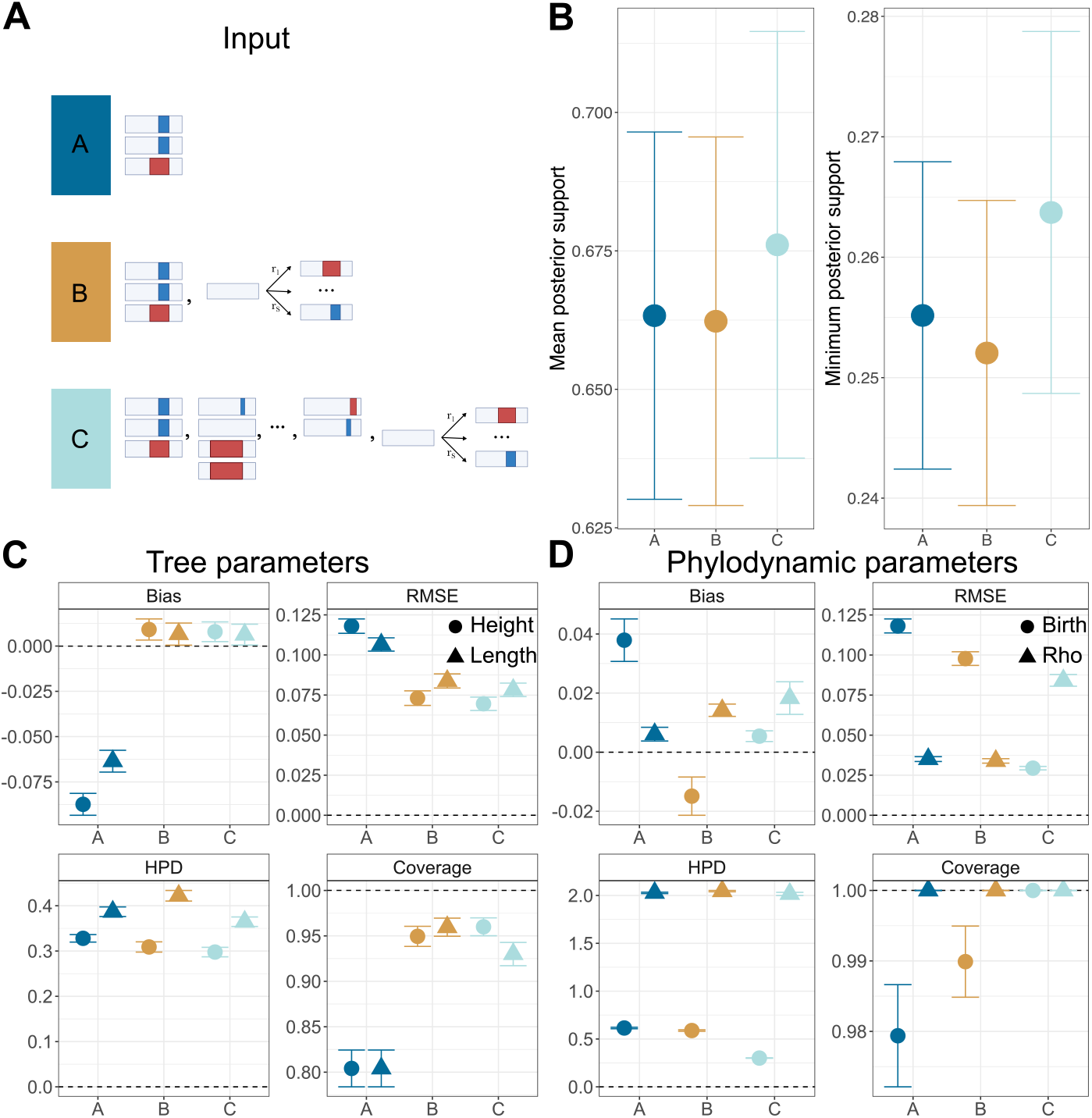
Inference performance when using different inputs. (A) Input description: A = barcode alignment; B = barcode alignment and scarring rates; C = 10 barcode alignments from experimental replicates and scarring rates. (B) Mean and minimum posterior support for the internal nodes of the true tree based on inputs as detailed in (A). Bias, RMSE, HPD width and coverage for the (C) tree height, tree length, (D) birth rate and sampling proportion based on input as detailed in (A). The error bars show the standard errors of the mean. The dashed line shows the best possible value for each metric.

Experimental replicates are a second source of information. Any variation between replicates should be due to the stochasticity of the process, not changes in the parameters describing it. We capture this scenario by using 10 alignments per inference instead of 1. Otherwise, we keep the inference settings as in the baseline, fix the scarring rates to the truth, infer a tree for each alignment and additionally share the cell division (birth rate) and death rate across trees (Fig. 3A, scenario C).

To compare the tree topology across different inputs, we assess the posterior probability of the true internal nodes with heights greater than 16 time units, which we call posterior support (Fig. 3B). In our simulation setup, nodes below 16 time units cannot be informed by the data and would only add noise to standard tree topology metrics. We compute the mean and minimum posterior support for each tree (i.e. the minimum or mean posterior probability across internal nodes) and average across trees for a given scenario. We find that the mean and minimum posterior support do not vary greatly (overlapping confidence intervals) between inputs. Nevertheless, for scenario C the average mean and minimum posterior support are highest.

We assess the accuracy of the tree and the phylodynamic parameters (Fig. 3C, D) using the bias and root mean square error (RMSE). Further we assess the precision and correctness using the 95% HPD width and the coverage.

For the tree parameters, the bias for scenarios B and C is 10 and the RMSE is 1.5 times smaller compared to scenario A. Differences between B and C are insignificant (confidence intervals overlapping). This indicates that known scarring rates greatly improve the accuracy of the tree parameters, while further adding experimental replicates does not add much information about the trees. Morerover, the HPD width for the tree height decreases slightly from 0.33± 0.01 in scenario A to 0.30 ± 0.01 in scenario C. For the tree length, the HPD width rises from 0.39 in A to 0.42 in B to 0.36 in C. The coverage for both tree parameters increases from ∼ 80% in scenario A to ∼ 95% in scenarios B and C. Hence, known scarring rates increase the correctness of the tree length and height but do not greatly increase the precision of their estimates.

For the phylodynamic parameters, we observe that the bias fluctuates for different parameters and scenarios. The RMSE for the birth rate decreases continually from 0.12 ± 0.004 in A to 0.1 ± 0.004 in B and 0.029 ± 0.001 in C. The RMSE for the death rate is similar for A and B (both 0.034 ± 0.0.002) while increased for scenario C (0.08 ± 0.004). However, the sum of the RMSEs for both parameters, decreases continually from A to C, indicating more accurate estimates for both parameters when more data is added. Additionally, while the HPD width for the death rate remains unchanged across inputs (2.0 ± 0.1), it becomes two times smaller for the birth rate in C (0.3 ± 0.002) compared to A and B (0.6 ± 0.01). Moreover, the coverage for the birth rate increases continually from A to C. Therefore, adding known scarring rates and experimental replicates leads to more accurate results, but only additionally adding experimental replicates results in more precise estimates.

### Benchmark and application on lineage tracing data

We benchmark TiDeTree on the intMEMOIR dataset (as available during the DREAM challenge) [14, 8] by comparing it against existing methods. The data is divided into a training and a test set. However, in most situations where lineage tracing is used, training data sets are not available. Hence, we use TiDeTree as an unsupervised method, i.e., we do not use the ground truth trees from the training set. Instead, we use all alignments within one inference, where we reconstruct a tree for each alignment while the scarring rates and phylodynamic parameters are shared across trees.

To estimate node heights, TiDeTree must assume a global editing rate, called a molecular clock rate in traditional phylogenetics. To test whether this assumption holds, we computed the proportion of edits at binned internal node intervals (Supp Fig. S2). The proportion of edits increases throughout the entire experimental period. Hence, we assume a molecular clock.

TiDeTree is among the top 3 methods in terms of topology inference (Fig. 4A). Here it is important to highlight that it outperforms many methods although it ignores the training set, i.e., uses less data. Moreover, in addition to a point estimate for the tree, we also report the posterior distribution of trees (Fig. 4B). This posterior can be visually inspected to assess which parts of the tree are well supported and which are uncertain. Further, for each tree posterior we constructed the 95% credible set, the smallest set of trees that make up 95% of the posterior. To construct this set, unique tree topologies with highest probability are continually added to the set, until their sum reaches 95%. The exact true tree topology is contained in the credible set in 31% of alignments. In fact, we noticed that some credible sets contain 10^4^ unique tree topologies, indicating little signal in the data to reliably favour one topology over another. Upon exclusion of credible sets with *>* 10^4^ trees, we recover the exact true tree topology within the credible set 74% of the time.

**Figure 4:**
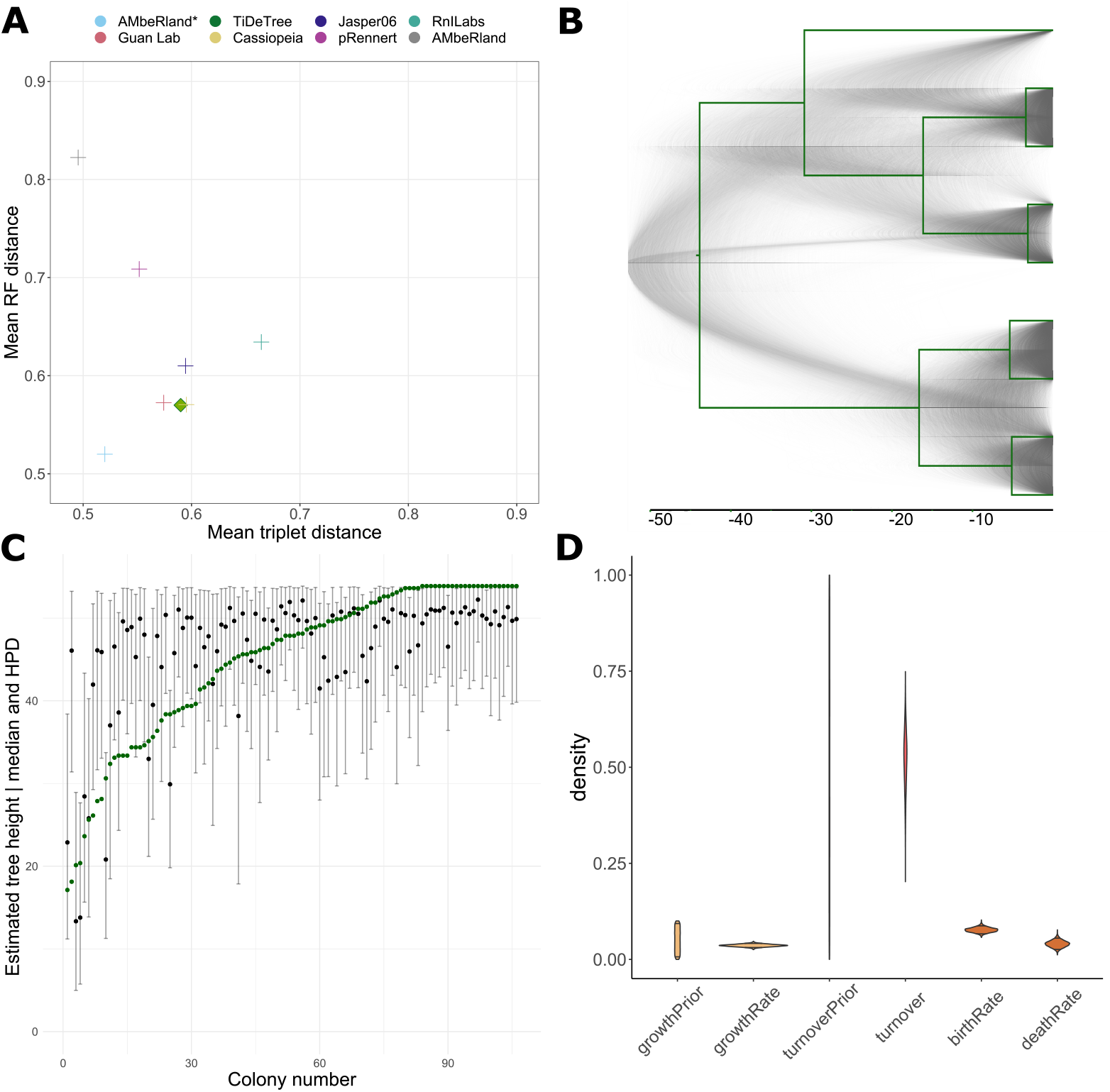
Application on lineage tracing data. (A) Topology benchmark ranking methods according to the Robinson-Foulds (RF) and triplet distance; TiDeTree is highlighted by a filled square. (B) Example tree posterior (grey) for colony s16 c4 allows to visually assess uncertainty in different parts of the tree. True tree is overlaid in green. (C) Estimated median tree height (black dots) and 95% HPD intervals for each dataset. The green dots show the true tree heights. (D) Estimated population dynamic parameters: prior and posterior distributions of the growth rate and turnover, posterior of the birth rate and death rate.

In a second analysis, we include additional model assumptions and prior information to get more reliable estimates of the node heights and the phylodynamic parameters.

First, the onset of editing is sometimes delayed (Supp Fig. S3. In principle, this could be due to the experimental setup, where the editing enzyme is induced at time 0. Then it must first be transcribed, translated and find its target DNA, before editing can commence. Therefore, we additionally sample the time of editing onset in the inference. Second, we use an independent dataset from [8], to place an informative prior on the scarring rate at each of the 10 target sites.

In this analysis the tree height is recovered in 82% of the time (Fig. 4, C). The median estimates and the 95% HPD intervals are 0.077 and [0.067, 0.087] per hour for the cell division rate and 0.026, [0.041, 0.055] per hour for the death rate. The cell division rate matches the reported number of ∼4 −5 generations over the 54*h* that were reported in the associated publication.

## Discussion

We introduced a time-dependent editing model and derive its likelihood calculation, enabling phylogenetic inference from lineage tracing data with independent targets. Our method, TiDeTree, estimates a time-scaled phylogeny, the scarring rates, and phylodynamic parameters within a Bayesian MCMC framework. Prior information and experimental replicates can easily be incorporated into the analysis to improve TiDeTree’s power. TiDeTree is available within the BEAST2 platform [6] allowing access to an ever-growing array of clock models, tree priors and other models. This enables users of TiDeTree immense flexibility to set up analyses that best fit their model systems. Frameworks for model selection by estimating the marginal likelihood [3, 20] and evaluating absolute model fit by posterior predictive simulation [11, 10] are also available and can be easily used in combination with our model. For instance, we assumed a strict clock model in our analyses, since we can assume that the editing rate does not vary over the short experimental time span. However, for longer time spans (e.g., during ontogenesis), the editing rate may vary over time. To account for this, relaxed clock models are readily available in BEAST2.

Several methods for inferring single-cell phylogenies were recently published. [14, 16, 13]. They have in common that they estimate the single best tree, while we derive posterior distributions of trees and all associated parameters of our model, allowing us to assess uncertainty directly. This will help developmental biologists to make more robust statements when using TiDeTree in their analyses. The method by [13] proposes a framework to assess temporal information in the data. Both their study and ours assume a constant editing rate (a so-called molecular clock assumption) in order to timescale the tree. However, whilst their method can only give a relative ordering along parallel lineages, TiDeTree estimates time-scaled trees, i.e. trees with branch lengths corresponding to the absolute time interval between cell divisions. Such time-scaled cell phylogenies will allow developmental biologists to address questions on the rates and timing of developmental events using lineage tracing data.

Current lineage tracing systems record edits at a limited number of sites, and for only a short time period compared to the entire experimental duration. Thus it is unclear how much information is contained about the developmental process; in particular how many population parameters can be concurrently estimated. In an ideal case two out of the three birth-death-sampling parameters can be obtained from a reconstructed tree [28]. We first evaluated the information content of the data based on our simulations. In our simulations, editing occurred for half of the experimental time span, and it appeared that only one parameter of a birth-death sampling model could be estimated from a single alignment. However, when we added additional information, e.g. by assuming scarring rates are known or by additionally adding experimental replicates, we could improve the accuracy, correctness and precision of the parameter estimates. Our analysis of an experimental lineage tracing dataset [8] supported this finding. Here, we could estimate two parameters, the cell division and death rate, by pooling them across 106 experimental replicates. These results highlight two possible routes to improving the signal for phylodynamic analyses. First, we can improve the signal contributed by individual trees, e.g. by increasing the number of targets (sites in the phylogenetic likelihood) or using repeatedly editing homing CRISPR barcode systems [17] to increase the editing time span. Second, we can include more experimental replicates and develop an approach through which they can inform complementary time spans. (Replicates 1–10 inform the first time span and replicates 10–20 inform second time span etc.).

Given the available data, statistical and computational challenges remain. First, on a technical level, a more efficient method for sampling subtrees with identical sequences is needed. Identical sequences arise because editing occurs only during a short time period and subsequent cell divisions lead to amplification of identical target sequences. However, we cannot simply collapse identical sequences to a single sequence because this would bias estimates of the tree generation process. Ideas might be co-opted from pathogen analyses that run into similar problems [5]. Further, specific clock models that are tailored to modelling editing through time as it occurs via lineage tracing could be developed.

Our contribution enables detailed analysis of developmental systems with different cell types. We view our methodology as a basis for Bayesian inference of single-cell time trees in conjunction with developmental parameters. By combining lineage tracing and single-cell RNAseq data, our approach allows to assess cell differentiation rates, as well as cell-type-specific division and apoptosis rates [27]. Such parameters quantify core processes of developmental biology. Thus, we see great promise that the advances in single cell lineage tracing technology combined with advances in single cell phylodynamic methodology will greatly enhance our understanding of developmental processes.

## Notes

### Competing Interest Statement

The authors have declared no competing interest.

