## Supplement for "TiDeTree: A Bayesian phylogenetic framework to estimate single-cell trees and population dynamic parameters from genetic lineage tracing data"

### Appendix

#### Transition probability matrix

For loss rate $l$ and scarring rates $s_i$ for $i = 1 : E$, define the following expressions:

$$r = l + \sum_{i=1}^{S} s_i \tag{5}$$

$$a_t = exp(-rt) \tag{6}$$

$$b_t = exp(-lt) \tag{7}$$

$$c_{it} = \frac{s_i(b_t - a_t)}{\sum_{i=1}^{S} s_i} \tag{8}$$

Then, the transition probability matrix outside of the scarring window, $P_{t:(t<t_1)|(t>t2)}$ is:

$$P_{t:(t<t_1)|(t>t2)} = \begin{pmatrix} b_t & 1-b_t & 0 & .. & 0 \\ 0 & 1 & 0 & .. & 0 \\ 0 & 1-b_t & b_t & .. & 0 \\ .. & .. & .. & .. & .. \\ 0 & 1-b_t & 0 & .. & b_t \end{pmatrix} \tag{9}$$

and the transition probability matrix for branches within the scarring window, $P_{t:(t_1 \leq t \leq t2)}$ is:

$$P_{t:(t_1 \leq t \leq t2)} = \begin{pmatrix} a_t & 1-b_t & c_{1t} & .. & c_{Et} \\ 0 & 1 & 0 & .. & 0 \\ 0 & 1-b_t & b_t & .. & 0 \\ .. & .. & .. & .. & .. \\ 0 & 1-b_t & 0 & .. & b_t \end{pmatrix} \tag{10}$$

These matrices can easily be verified by showing that the following equality holds for each one of them:

$$\frac{d}{dt} P_t = Q \times P_t \tag{11}$$

*Proof.*

$$\frac{d}{dt}P_{t:(t<t_1)|(t>t2)} = Q \times P_{t:(t<t_1)|(t>t_2)}$$

$$\begin{pmatrix} -l\,exp[-lt] & l\,exp[-lt] & 0 \\ 0 & 0 & 0 \\ 0 & l\,exp[-lt] & -l\,exp[-lt] \end{pmatrix} = \begin{pmatrix} -l & l & 0 \\ 0 & 0 & 0 \\ 0 & l & -l \end{pmatrix} \times \begin{pmatrix} b_t & 1-b_t & 0 \\ 0 & 1 & 0 \\ 0 & 1-b_t & b_t \end{pmatrix}$$

$$\begin{pmatrix} -l\,b_t & l\,b_t & 0 \\ 0 & 0 & 0 \\ 0 & l\,b_t & -l\,b_t \end{pmatrix} = \begin{pmatrix} -l\,b_t & l\,b_t & 0 \\ 0 & 0 & 0 \\ 0 & l\,b_t & -l\,b_t \end{pmatrix}$$

□

*Proof.*

$$\frac{d}{dt}P_{t:(t_1\le t\le t2)} =$$

$$\begin{pmatrix} -r\,exp[-rt] & l\,exp[-lt] & \frac{s_1(r\,exp[-rt]-l\,exp[-lt])}{\sum_i s_i} & \dots & \frac{s_S(r\,exp[-rt]-l\,exp[-lt])}{\sum_i s_i} \\ 0 & 0 & 0 & \dots & 0 \\ 0 & l\,exp[-lt] & -l\,exp[-lt] & \dots & 0 \\ \dots & \dots & \dots & \dots & \dots \\ 0 & l\,exp[lt] & 0 & \dots & -l\,exp[-lt] \end{pmatrix} =$$

$$\begin{pmatrix} -r\,a_t & l\,b_t & \frac{s_1(r\,a_t-l\,b_t)}{\sum_i s_i} & \dots & \frac{s_S(r\,a_t-l\,b_t)}{\sum_i s_i} \\ 0 & 0 & 0 & \dots & 0 \\ 0 & l\,b_t & -l\,b_t & \dots & 0 \\ \dots & \dots & \dots & \dots & \dots \\ 0 & l\,b_t & 0 & \dots & -l\,b_t \end{pmatrix} =$$

$$\begin{pmatrix} -r & l & s_1 & \dots & s_S \\ 0 & 0 & 0 & \dots & 0 \\ 0 & l & -l & \dots & 0 \\ \dots & \dots & \dots & \dots & \dots \\ 0 & l & 0 & \dots & -l \end{pmatrix} \times \begin{pmatrix} a_t & 1-b_t & c_{1t} & .. & c_{St} \\ 0 & 1 & 0 & .. & 0 \\ 0 & 1-b_t & b_t & .. & 0 \\ .. & .. & .. & .. & .. \\ 0 & 1-b_t & 0 & .. & b_t \end{pmatrix} =$$

$$Q \times P_{t:(t_1<t<t_2)}$$

□

#### Implementation

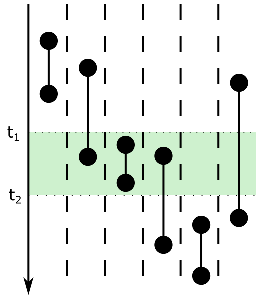


Figure S1: **Branch types** For branch types 1, 3, 5, the transition probability can be calculated as usual. For branch types 2 and 4, the branch has to be cut once, at $t_1$ and at $t_2$, respectively. For branch type 6 the branch has to be cut twice at both $t_1$ and $t_2$.

#### Validation

| Parameter | Coverage | 95% HPD width | RMSE | Bias |
|---|---|---|---|---|
| treeHeight | 0.94 | 1.32 | 0.35 | 0.22 |
| treeLength | 0.93 | 1.50 | 0.39 | 0.22 |
| birthRate | 0.96 | 0.25 | 0.05 | 0.00 |
| deathRate | 1.00 | 0.00 | 0.00 | 0.00 |
| rho | 0.96 | 1.63 | 0.36 | 0.15 |
| scarringRate.1 | 0.97 | 0.02 | 0.98 | -0.98 |
| scarringRate.2 | 0.98 | 0.02 | 0.98 | -0.98 |
| scarringRate.3 | 0.97 | 0.02 | 0.98 | -0.98 |
| scarringRate.4 | 0.98 | 0.02 | 0.98 | -0.98 |
| scarringRate.5 | 0.98 | 0.02 | 0.98 | -0.98 |
| scarringRate.6 | 0.97 | 0.02 | 0.98 | -0.98 |
| scarringRate.7 | 0.99 | 0.02 | 0.99 | -0.99 |
| scarringRate.8 | 0.97 | 0.02 | 0.98 | -0.98 |
| scarringRate.9 | 0.98 | 0.02 | 0.99 | -0.99 |
| scarringRate.10 | 0.98 | 0.02 | 0.99 | -0.99 |
| scarringRate.11 | 0.98 | 0.02 | 0.99 | -0.99 |
| scarringRate.12 | 0.98 | 0.02 | 0.99 | -0.99 |
| scarringRate.13 | 0.97 | 0.02 | 0.98 | -0.98 |
| scarringRate.14 | 0.98 | 0.02 | 0.98 | -0.98 |
| scarringRate.15 | 0.98 | 0.02 | 0.98 | -0.98 |
| scarringRate.16 | 0.98 | 0.02 | 0.99 | -0.99 |
| scarringRate.17 | 0.97 | 0.02 | 0.98 | -0.98 |
| scarringRate.18 | 0.98 | 0.02 | 0.99 | -0.99 |
| scarringRate.19 | 0.98 | 0.02 | 0.99 | -0.99 |
| scarringRate.20 | 0.98 | 0.02 | 0.99 | -0.99 |
| scarringRate.21 | 0.97 | 0.02 | 0.98 | -0.98 |
| scarringRate.22 | 0.98 | 0.02 | 0.98 | -0.98 |
| scarringRate.23 | 0.98 | 0.02 | 0.98 | -0.98 |
| scarringRate.24 | 0.98 | 0.02 | 0.99 | -0.99 |
| scarringRate.25 | 0.98 | 0.02 | 0.98 | -0.98 |
| scarringRate.26 | 0.98 | 0.02 | 0.99 | -0.99 |
| scarringRate.27 | 0.98 | 0.02 | 0.99 | -0.99 |
| scarringRate.28 | 0.98 | 0.02 | 0.99 | -0.99 |
| scarringRate.29 | 0.98 | 0.02 | 0.99 | -0.99 |
| scarringRate.30 | 0.98 | 0.02 | 0.99 | -0.99 |
| scarringRate.31 | 0.98 | 0.02 | 0.99 | -0.99 |
| scarringRate.32 | 0.98 | 0.02 | 0.99 | -0.99 |
| scarringRate.33 | 0.98 | 0.02 | 0.99 | -0.99 |
| scarringRate.34 | 0.99 | 0.02 | 0.98 | -0.98 |
| scarringRate.35 | 0.98 | 0.02 | 0.99 | -0.99 |
| scarringRate.36 | 0.98 | 0.02 | 0.99 | -0.99 |
| scarringRate.37 | 0.99 | 0.02 | 0.99 | -0.99 |

| | | | | |
|---|---|---|---|---|
| scarringRate.38 | 0.98 | 0.02 | 0.99 | -0.99 |
| scarringRate.39 | 0.98 | 0.02 | 0.99 | -0.99 |
| scarringRate.40 | 0.97 | 0.02 | 0.98 | -0.98 |
| scarringRate.41 | 0.97 | 0.02 | 0.98 | -0.98 |
| scarringRate.42 | 0.98 | 0.02 | 0.99 | -0.99 |
| scarringRate.43 | 0.98 | 0.02 | 0.99 | -0.99 |
| scarringRate.44 | 0.98 | 0.02 | 0.98 | -0.98 |
| scarringRate.45 | 0.97 | 0.02 | 0.99 | -0.99 |
| scarringRate.46 | 0.98 | 0.02 | 0.98 | -0.98 |
| scarringRate.47 | 0.97 | 0.02 | 0.98 | -0.98 |
| scarringRate.48 | 0.97 | 0.01 | 0.99 | -0.99 |
| scarringRate.49 | 0.98 | 0.02 | 0.98 | -0.98 |

Table S1: Metrics for well-calibrated simulations.

#### Including additional info

| Scarring rate index | Value |
|---|---|
| 1 | 0.38 |
| 2 | 0.59 |
| 3 | 0.07 |
| 4 | 0.07 |
| 5 | 0.22 |
| 6 | 1.45 |
| 7 | 0.61 |
| 8 | 0.27 |
| 9 | 0.48 |
| 10 | 0.07 |
| 11 | 0.7 |
| 12 | 0.38 |
| 13 | 0.62 |
| 14 | 2.21 |
| 15 | 0.53 |
| 16 | 0.52 |
| 17 | 0.94 |
| 18 | 0.33 |
| 19 | 0.17 |
| 20 | 0.29 |
| 21 | 1.18 |
| 22 | 0.32 |
| 23 | 0.15 |
| 24 | 0.28 |
| 25 | 0.05 |
| 26 | 0.03 |
| 27 | 0.29 |
| 28 | 1.98 |
| 29 | 0.59 |
| 30 | 0.5 |
| 31 | 0.72 |
| 32 | 0.02 |
| 33 | 0.16 |
| 34 | 0.66 |
| 35 | 0.1 |
| 36 | 0.51 |
| 37 | 0.15 |
| 38 | 0.36 |
| 39 | 0.38 |
| 40 | 0.12 |
| 41 | 0.54 |
| 42 | 0.51 |

| | |
|---|---|
| 43 | 0.65 |
| 44 | 0.63 |
| 45 | 0.28 |
| 46 | 0.15 |
| 47 | 0.65 |
| 48 | 0.5 |
| 49 | 0.26 |
| 50 | 1 |

Table S2: Scarring rates used in simulation and during the inference for scenarios B and C.

#### intMemoir

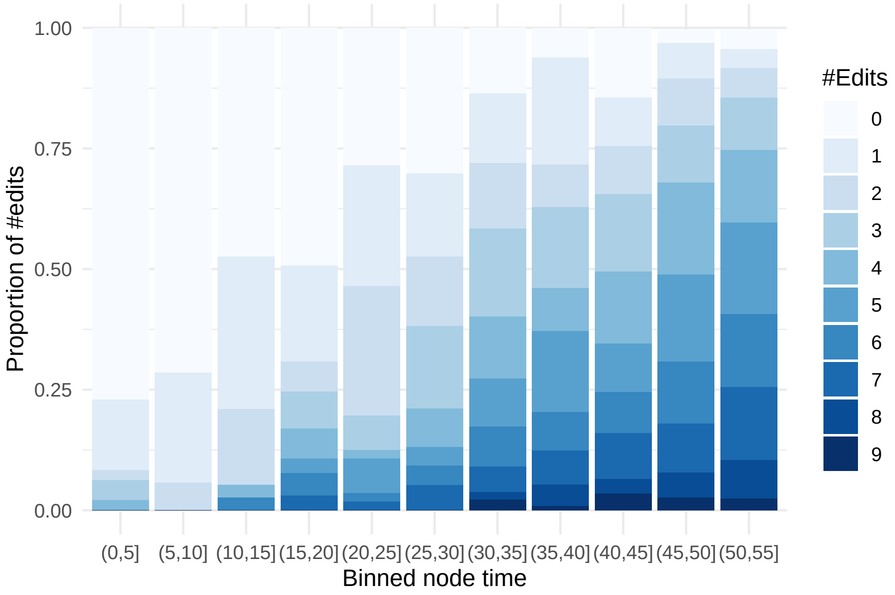


Figure S2: **Temporal signal in intMEMOIR data.** Proportion of edits against binned node time increases through time. That means, the more time passes during the experiment, the more barcodes are scarred on average. Hence, a molecular clock model should hold for the accumulation of scars.

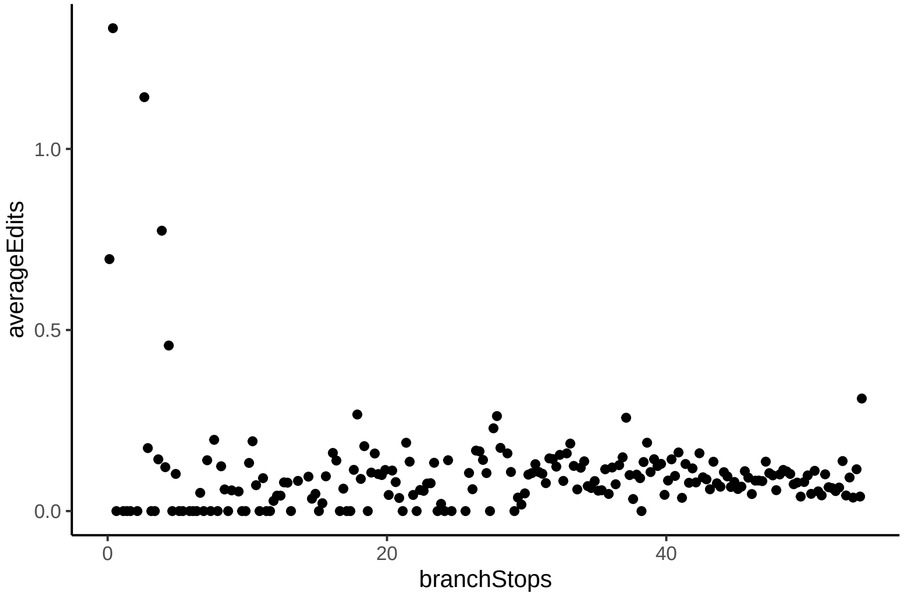


Figure S3: **Editing start.** The average number of edits for all branches, that stop at time x. We observe that at the beginning of the experiment, there are still a lot of branches on which no edits occur. This could be caused by a delayed editing start as the transposase expression is only induced at time 0. If we let the editing start vary in our inference, we recover the true tree height better.
